# Making Moral Decisions With Artificial Agents As Advisors. An *f*NIRS Study

**DOI:** 10.1101/2024.03.05.583335

**Authors:** Eve Florianne Fabre, Damien Mouratille, Vincent Bonnemains, Grazia Pia Palmiotti, Mickael Causse

**Author notes:** Correspondence should be addressed to Eve F. Fabre. Author Note The present research was funded by a postdoctoral grant awarded by the AXA research fund to Eve F. Fabre. We would like to thank Pr. Frédéric Dehais and Dr. Sébastien Scanella for their help with the *f*NIRS device.

## Abstract

Artificial Intelligence (AI) is on the verge of impacting every domain of our lives. It is increasingly being used as an advisor to assist in making decisions. The present study aimed at investigating the influence of moral arguments provided by AI-advisors (i.e., decision aid tool) on human moral decision-making and the associated neural correlates. Participants were presented with sacrificial moral dilemmas and had to make moral decisions either by themselves (i.e., baseline run) or with AI-advisors that provided utilitarian or deontological arguments (i.e., AI-advised run), while their brain activity was measured using an *f*NIRS device. Overall, AI-advisors significantly influenced participants. Longer response times and a decrease in right dorsolateral prefrontal cortex activity were observed in response to deontological arguments than to utilitarian arguments. Being provided with deontological arguments by machines appears to have led to a decreased appraisal of the affective response to the dilemmas. This resulted in a reduced level of utilitarianism, supposedly in an attempt to avoid behaving in a less cold-blooded way than machines and preserve their (self-)image. Taken together, these results suggest that motivational power can led to a voluntary up- and down- regulation of affective processes along moral decision-making.

## 1. INTRODUCTION

### 1.1. The Revolution of Artificial Intelligence

The revolution of Artificial Intelligence (AI) is impacting almost every domain of our life. At home, social chatbots (<u>Henkel et al., 2020</u>; <u>Pentina et al., 2023</u>) are being increasingly used as personal assistants (e.g., Alexa, Siri, Chat GPT, Google Assistant), friendship companions (e.g., Replika, Anima, Kajiwoto, Microsoft XiaoIce) or even relational agents for (mental) healthcare (e.g., Woebot; Wysa; Youper; <u>Stade et al., 2023</u>). At work, intelligent decision support systems are now being used as advisors in healthcare, finance, and the military to name a few (<u>Černevičienė & Kabašinskas, 2022</u>; <u>Macey-Dare, 2023</u>; <u>Shortliffe, & Sepúlveda, 2018</u>; <u>Wasilow & Thorpe, 2019</u>). These systems allow the fusion and analysis of large amounts of data to reveal complex relationships, provide recommendations to workers and help them make faster and more informed decisions compared to when the data is analyzed manually (<u>Eom & Kim, 2006</u>). The increasing use of AI-agents as advisors, cooperators and sometimes even delegates, raises the question of whether and to what extent do AI decision aids (i.e., AI advisors) modulate human beings’ judgment and decision-making (e.g., <u>Collart et al., 2015</u>; <u>Crandall et al., 2018</u>; <u>Grgić-Hlača, et al., 2019</u>; <u>Hanson et al., 2024</u>; <u>Köbis et al., 2021; Ladak et al., 2024</u>). The present study aimed at investigating this question, and more specifically whether moral advice provided by AI-advisors can influence human beings’ moral decisions.

### 1.2. Using Moral Dilemmas to Study Moral Decision-Making

Moral dilemmas, particularly sacrificial ones, have been extensively used to explore moral judgment and decision-making, alongside the affective and cognitive processes underlining these judgments and decisions (e.g., <u>Christensen & Gomila, 2012</u>; <u>Greene et al., 2001</u>, <u>2015</u>). When confronted with a sacrificial moral dilemma (<u>Patil et al., 2021</u>), one usually has to decide between a *utilitarian option*, which involves sacrificing the few to save the many (<u>Mill, 1998</u>), and a *deontological option*, which entails avoiding causing harm to individuals unrelated to the situation and refraining from intervening (<u>Kant, 2013</u>). Individuals’ moral decisions tend to vary depending on the dilemma, as reflected by the disparity in moral preferences commonly observed between dilemmas such as the trolley dilemma and the footbridge dilemma (<u>Christensen & Gomila, 2012</u>). In both dilemmas, one must envision that a trolley is hurtling at full speed along a track, where five workers stand, and is about to strike and kill these workers. In the trolley dilemma, the sole means of saving the workers is to pull a lever, which would divert the trolley onto a sidetrack where only one worker is present (<u>Foot, 1978</u>; <u>Thomson, 1985</u>). Conversely, in the footbridge dilemma, saving the workers involves pushing an overweight individual off the footbridge and into the path of the oncoming trolley (<u>Thomson, 1976)</u>. Despite the fact that the two options to save the workers adhere to a utilitarian principle of sacrificing one life to save five, the greater majority of people opts for pulling the lever in the trolley dilemma but refuse to push the overweight person off the footbridge.

Exploring the reasons behind the preferences for utilitarian versus deontological options has been a central focus of numerous studies in the field of moral neuroscience (<u>Casebeer, 2003; Greene et al, 2001</u>, <u>2004</u>, <u>2015</u>; <u>Moll et al., 2005</u>). Under some circumstances, the utilitarian option can trigger a strong negative affective reaction, reflected by the activity of the ventromedial prefrontal cortex (vmPFC; <u>Moll & de Oliveira-Souza, 2007</u>) and the temporo-parietal junction (TPJ). This strong negative reaction occurs for instance when it involves killing someone as an intended means for the greater good like in the footbridge dilemma (<u>Foot, 1978</u>), and usually results in a preference for the deontological option. In other circumstances, such as when the death of an individual unrelated to the situation is a foreseen but unintended consequence (e.g., <u>Sarlo et al., 2012</u>), the utilitarian option triggers an increased activity of brain regions associated with working memory and problem solving, such as the dorsolateral prefrontal cortex (DLPFC; <u>Moll et al., 2005</u>), reflecting the greater cognitive effort made to control the negative affective response. In this case, the utilitarian option is more likely to be preferred (<u>Greene, 2015</u>; <u>Tassy et al., 2012</u>; <u>Zheng et al., 2018</u>). Overall, this literature aligns with the dual process theory of moral judgment, which postulates that moral decisions result from two competing processing systems: a fast, automatic, and emotional System 1 and a slow, controlled and cognitively costly System 2 (<u>Greene, 2015</u>).

### 1.3. The Influence of Human and Artificial Advisors

Human beings are organized in social groups, wherein they constantly exchange knowledge (<u>Gariépy et al., 2014</u>) and seek advice (e.g., <u>Polman & Ruttan, 2022</u>). As a result, they exhibit a heightened sensitivity to the influence of their peers (<u>Cialdini & Goldstein, 2004; Yu et al., 2021</u>). They tend to align their judgments and behaviors with those of their peers (e.g., <u>Braams et al., 2019</u>; <u>Chung et al., 2020</u>; <u>Fabre et al., 2022</u>) and being swayed by their advice (e.g., <u>Bonaccio & Dalal, 2006</u>; <u>Leong & Zaki, 2018</u>; <u>McCoy & Natsuaki, 2015</u>). The extent of an advisor’s influence is highly dependent on the advisee’s trust, the advisors’ expertise and reliability (<u>Bonaccio & Dalal, 2006;</u> for a meta-analysis see <u>Bailey et al., 2023</u>). Overall, neuroscience studies have shown an increased activity in the lateral orbitofrontal cortex and the vmPFC in response to advice that contradicts one’s own opinions, and greater activity in the ventral striatum and the anterior cingulate cortex when the difference in opinion with the advisor is low (e.g., <u>Biele et al., 2011</u>; <u>Campbell-Meiklejohn et al., 2010</u>; <u>Izuma, 2017</u>; <u>Meshi et al., 2012</u>). The activity of these brain regions is known to increase in response to respectively negative and positive outcomes, suggesting that sharing similar opinions is socially rewarding for advisees (<u>Izuma, 2017</u>). Interestingly, the activity of left lateral orbitofrontal cortex and the DLPFC was found to be positively correlated to the influence of the advice (<u>Meshi et al., 2012; Zhang & Gläscher, 2020)</u>.

In the last decades, an important number of studies have demonstrated that non-human agents can also have a significant influence on humans (for an overview, see <u>Hertz & Wiese, 2019</u>). The influence of non-human agents’ advice was found to vary as a function of the non-human advisors’ perceived expertise (as for human advisors), but also depending on their similarity to humans in terms of appearance, empathy or non-verbal cues (e.g., <u>Benitez et al, 2017; Chidambaran et al., 2012; Goodyear et al., 2016</u>, <u>2017</u>). Part of this literature focused specifically on the influence of non-human agents on moral judgment and/or decision-making (<u>Jackson & Williams, 2019</u>; <u>Köbis et al., 2021</u>; <u>Leib et al., 2024; Straßmann et al., 2020</u>). Some studies have shown that artificial advisors can have a corruptible impact on humans, increasing for instance their tendency to break ethical rules (e.g., cheat or inflict harm to another individual) for their own profit (e.g., monetary gain; <u>Köbis et al., 2021; Sandoval et al., 2016</u>). While there is a substantial body of literature examining how humans judge the decisions of machines in sacrificial moral dilemma situations (e.g., <u>Awad et al., 2018</u>, <u>2019</u>; <u>Bonnefon et al., 2016</u>; <u>Malle et al., 2015</u>), the influence of non-human advisors on human decision-making in these specific situations has received very limited attention (<u>Hanson et al., 2024</u>; <u>Straßmann et al., 2020)</u>.

### 1.4. The present study

The aim of the present study was to investigate the influence of moral arguments provided by AI-advisors, acting as decision aid tool, on human moral decisions and the associated neural correlates. Participants had to decide on sacrificial moral dilemmas (<u>Patil et al., 2021</u>) that were either categorized as *utilitarian dilemmas* (e.g., the trolley dilemma; <u>Foot, 1978</u>; <u>Thompson, 1985</u>) or *deontological dilemmas* (e.g., the footbridge dilemma; <u>Thomson, 1976</u>), depending on the general preference for the utilitarian option or the deontological option (i.e., rating study). In the baseline run, participants made their decisions by themselves, while in the AI-advised run, AI-advisors had already pre-selected one of the two options and provided participants with an argument to justify these pre-decisions. In the AI-advised run, participants also had the possibility to delegate the execution of the decisions to the AI-advisors by not answering for more than 15s. In this run, participants’ prefrontal cortex activity was assessed using an *f*NIRS system (<u>Balconi & Fronda, 2020</u>; <u>Dashtestani et al., 2018</u>, <u>2019</u>; <u>Lee & Yun, 2017</u>; <u>Strait & Scheutz, 2014</u>).

Behavioral and brain activity data were first analyzed in terms of change of mind rate (compared to the baseline), as a function of the argument valence (i.e., validating versus contradicting the decision made in the baseline run). The data were also further analyzed as a function of the type of the dilemma (i.e., utilitarian/deontological) and the type of argument (i.e., utilitarian/deontological). Overall, we predicted to observe a greater proportion of utilitarian decisions in response to utilitarian dilemmas, and a greater proportion of deontological decisions in response to deontological dilemmas. Regarding brain activity, we predicted to observe a greater activity of the DLPFC in response to deontological dilemmas than to utilitarian dilemmas (<u>Greene et al., 2001</u>, <u>2004</u>), reflecting the greater cognitive effort made to control the initial negative affective response to deontological dilemmas (<u>Jeurissen et al., 2014</u>; <u>Lee & Yun, 2017</u>). Regarding the influence of AI-advisors, we predicted to observe 1) a greater change of mind rate in response to contradicting arguments than to validating arguments and 2) a significant increase in utilitarian decisions in response to utilitarian arguments and/or a significant decrease in utilitarian decisions in response to deontological arguments in the AI-advised run (compared to the baseline run), with a potential interaction effect with the type of dilemma. We predicted that the influence of the AI-advisors would ensue from participants’ motivation to engage in a greater cognitive effort to process contradicting arguments, which might be reflected by the greater activity of the DLPFC in response to contradicting arguments than to validating arguments (<u>Jeurissen et al., 2014</u>; <u>Lee & Yun, 2017</u>). We did not have specific predictions regarding participants’ decisions to use AI advisors as delegates (i.e., to outsource the execution of their moral decisions) due to conflicting findings in previous studies. Some studies have found that participants retained control of action execution (<u>Bigman & Gray, 2018</u>; <u>Castelo et al., 2019</u>; <u>Dietvorst et al., 2015</u>, <u>2018</u>; <u>Gogoll & Uhl, 2018</u>), while others found that they can cede their control authority to non-human agents (<u>Gombolay et al., 2015</u>; <u>Leyer & Schneider, 2019</u>; <u>Robinette et al., 2016</u>).

## 2. METHODS

### 2.1. Participants

A review of the literature for sample size estimation c performed with the Pingouin package (v.0.5.3) in Python (v3.11) c revealed the necessity to record at least 8 participants to observe both the effect of the dilemma type on *f*NIRS (<u>Lee & Yun, 2017</u>) and the effect of advice type on performances (<u>Goodyear et al., 2016</u>) with a power (1 - β) = 95% and a significance level α = 5%. In addition, the sample size estimation for the effect of the interaction between dilemma type and argument type could not be computed since this has never been done with *f*NIRS to our knowledge. The closest result can be obtained by computing the interaction between dilemma type and response type on RMI data (<u>Dashtestani et al., 2018</u>) which resulted in a sample size of ∼32 participants required to observe this effect with a power of 95% and a significance of 5%. 40 French participants (14 females, *Mage* = 28 years old) participated in the study. All were right-handed and had normal or corrected-to-normal vision. None of the participants reported a history of prior neurological disorder. They received no compensation for their participation.

### 2.2. Ethical concerns

All participants were informed of their rights and gave written informed consent for participation in the study. This study was carried out in accordance with both the Declaration of Helsinki and the recommendations of the Research Ethics Committee of the University of XXX in XXX (XXX no. 2017-045).

### 2.3. Materials

#### 2.3.1. Selection of the dilemmas and the arguments

Two rating studies were conducted to select and balance the experimental materials. A first rating study was conducted to select an equal number of utilitarian dilemmas and deontological dilemmas, the two groups of dilemmas being balanced in terms of choice ratio (e.g., a utilitarian dilemma with a utilitarian choice rate of 80% was balanced with a deontological dilemma with a deontological choice rate of 80%). A second rating study was conducted to select one utilitarian argument and one deontological argument per dilemma (i.e., respectively arguing that *“it is necessary to sacrifice the few to save the many”* versus that *“unrelated people should not be sacrificed to save others”*) c the two classes of arguments being balanced in terms of persuasiveness. The rating studies are described in detail in the supplementary materials. The 18 dilemmas and the 36 arguments that were used in the present study are presented in Table 1S.

#### 2.3.2. Experimental apparatus

The experiment was designed to record a hemodynamic neuronal response using *f*NIRS. The *f*NIRS signals were acquired by the NIRScout device manufactured by the NIRx Company (Germany). The NIRScout system has a 7.8125 Hz resolution and contains eight sources and seven detectors placed on the subject’s scalp as shown in Figure 1. Eight optodes were placed close to the sources (i.e., less than 1 cm) to create eight short-channels. Each “source–detector” pair was placed close enough to each other (∼ 3 cm) to form a *f*NIRS channel, generating 16 channels. Three regions of interest (ROI) were defined: 1) the rostral-lateral prefrontal area (RLPFC) composed by the S4-D3, S4-D4, S3-D3, S5-D4, S5-D5 and S6-D5 source-detector pairs, 2) the right dorsolateral prefrontal cortex (right-DLPFC) composed by S1-D1, S2-D1, S2-D2, S3-D2 and S3-D1 source-detector pairs and 3) the left dorsolateral prefrontal cortex (left-DLPFC) composed by S6-D6, S7-D6, S7-D7, S6-D7 and S8-D6 source-detector pairs (see Figure 1).

**Figure 1.**
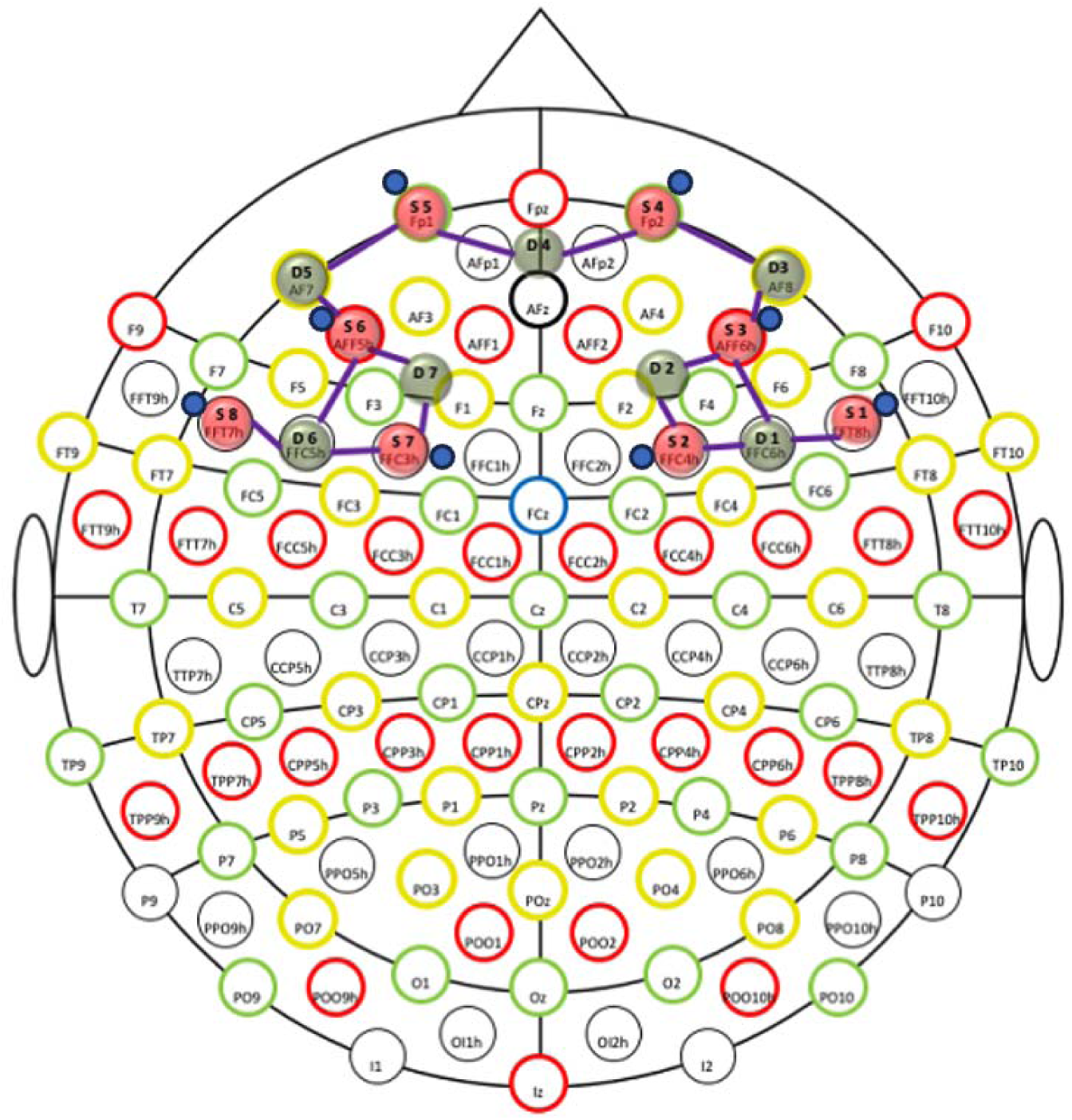
Illustration of the *f*NIRS montage in international 10-10 coordinate space. Montage with 8 × 9 frontal source-detector pairs. Sources are colored in red, detectors are colored in green, optodes for short-channels are colored in blue, and channels are indicated by purple lines.

### 2.4. Task

The experiment was divided into two runs. In the baseline run, participants were presented with 18 sacrificial moral dilemmas (i.e., nine utilitarian and nine deontological), each time with two possible options: one utilitarian and one deontological. Participants read the dilemma and then pressed the spacebar to make it disappear. After a 1-second blank screen, the two options were displayed side by side. Each type of option appeared half of the time on the left side of the screen and half of the time on the right side. Participants had to choose one of the two options by pressing a key on an AZERTY keyboard: the “*a*” key for the option on the left or the “*p*” key for the option on the right (see Figure 2).

**Figure 2.**
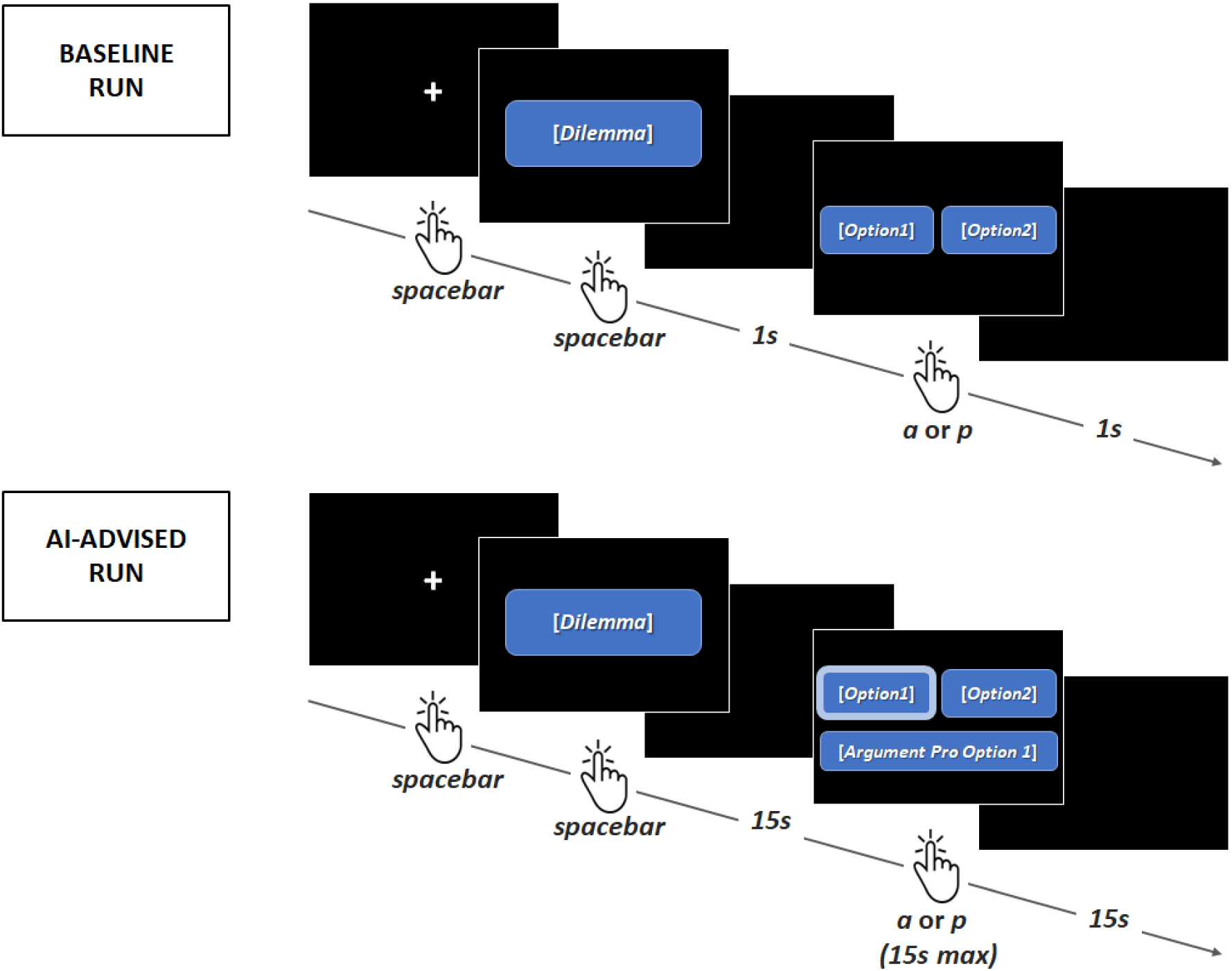
Illustrations of a trial in 1) the baseline run (top) and 2) the AI-advised run (bottom).

In the AI-advised run, participants were presented with the same sacrificial dilemmas as in the baseline run, but with different AI-advisors serving as moral advisors (i.e., a decision-aid tool based on artificial intelligence). Each dilemma was presented twice: once with an AI-advisor that had preselected the utilitarian option and provided a utilitarian argument and once with an AI-advisor that had preselected the deontological option and provided a deontological argument. As in the baseline run, participants read the dilemma and then pressed the spacebar to make it disappear. After a 15s blank screen, participants were presented with the two options (the option preselected by the AI-advisor was circled in light blue) and the argument justifying the pre-decision of the artificial advisor. In this run too, participants had to choose one of the two options by pressing a key on an AZERTY keyboard: the “*a*” key for the option on the left or the “*p*” key for the option on the right. They also had the possibility to remain inactive for 15s and let the AI-advisor execute its pre-decision (i.e., delegation). Participants were given no information regarding the characteristics of the AI-advisors (e.g., reliability, design).

### 2.5. Procedure

After signing the informed consent form, participants were asked to enter the experimental room and read the instructions for the experiment. They were informed that in the first phase of the experiment they would be presented with different sacrificial dilemmas and their task was to choose between two options. There was no time limit for them to give their answer. It was emphasized that there were no right or wrong answers, and they should make their decisions based on their personal preferences. Participants were invited to ask any questions before the experiment began. They then sat in a comfortable seat in front of the 1920 x 1080 computer screen and performed the baseline run. At the end of the baseline run, the NIRScout system was placed on their heads. They were equipped with the NIRScap, to which 23 optodes were attached. The NIRX system was then calibrated. Pre-tests prompted us to record participants’ brain activity only during the AI-advised run to prevent them from experiencing headaches triggered by the *f*NIRS system. Concurrently to the installation of the *f*NIRS system, participants were given the written instructions for the second phase of the experiment (i.e., AI-advised run). They were informed that in the second phase of the experiment, their task would be the same as in the first phase, but this time they would receive advice from different AI-advisors. It was explained that the arguments presented had been pre-generated by various AIs, each trained on distinct sets of human moral decision-making arguments. Participants were asked to imagine that they were making decisions with the assistance of these AI-advisors, which could automatically analyze each situation and provide guidance. Once ready, participants performed the AI-advised run. At the end of the latter, the experimenter removed the NIRScout system from the participants’ head. Participants were then asked to fill the French version of the Interpersonal Reactivity Index (IRI; composed of a empathic concern, a perspective taking, a personal distress and a fantasy subscale; <u>Davis, 1980</u>; <u>Gilet et al., 2013</u>), the Gudjonsson’s compliance scale that we translated in French (<u>Gudjonsson, 1989</u>) and the French version of the Negative Attitudes towards Robots Scale (NARS; composed of an interaction, a social and an emotion subscale; <u>Dinet & Vivian, 2015</u>; <u>Nomura et al., 2006</u>). When filling the NARS and the Gudjonsson’s compliance scale, participants had to indicate their level of agreement with the statements on respectively a 9-point Likert scale (from – 4 = *totally disagree* to 4 = *totally agree*) and a 7-point Likert scale (from 1 = *totally disagree* to 7 = *totally agree*). When filling the IRI, participants had to indicate the extent to which the statements described them on a 7-point Likert scale (from 1 = *does not describe me at all* to 7 = *describes me perfectly*). After completing the questionnaires, participants were debriefed. They were asked about their general impressions of the experiment and their thoughts on the possibility of delegating the execution of actions to AI-advisors. They were invited to ask any questions they had. Finally, they were thanked for their participation and left the experimental room.

### 2.6. Data measures

#### 2.6.1. Behavioral measures

##### Decisions

Participants’ decisions were assessed in terms of utilitarian decisions. Delegations (see below) were counted as utilitarian decisions when the AI-advisors had preselected the utilitarian option.

##### Changes of mind

To further analyze participants’ decisions, the decisions made in the AI-advised run were compared to those made in the baseline run (for each dilemma), a measure referred to as changes of mind in the present study. Changes of mind were assessed in terms of 1) changes of mind from the deontological option to the utilitarian option, 2) changes of mind from the utilitarian option to the deontological option and 3) indiscriminate changes of mind. Delegations were taken into account based on the option preselected by the AI-advisors.

##### Delegations versus active decisions

A delegation was defined as an absence of participants’ response for more than 15s that triggered the execution of the AI-advisors’ pre-decision. They were assessed in terms of active decisions.

##### Response times

In the baseline run, response times were defined as the time period between the onset of the options and the beginning of the participants’ responses. In the AI-advised run, they were defined as the time period between the onset of the options and the AI-advisors’ arguments and the beginning of the participants’ responses. The response times above or below two standard deviations around the mean and those associated with delegations were eliminated (<u>Ratcliff, 1993</u>). Response times were assessed in terms of mean response times.

#### 2.6.2. *f*NIRS measures

*f*NIRS data was preprocessed by using MNE-Python (<u>Gramfort et al., 2013</u>), its extension MNE-NIRS (<u>Luke et al., 2021</u>) and following the guidelines from <u>Yücel and colleagues (2021</u>). First, raw data were converted into optical density data. Then, a channel pruning was applied using the scalp-coupling index (SCI) for each channel. SCI is an indicator of the quality of the connection between the optodes and the scalp, which looks for the presence of a prominent synchronous signal in the frequency range of cardiac signals across the photo-detected signals (<u>Pollonini et al., 2014</u>). Channels with a scalp-coupling index below 0.75 were marked as bad channels (less than 10%) and were interpolated with the nearest channel providing good data quality. Visual checking of data was also performed. In order to remove baseline shift and spike artifacts, temporal derivative distribution repair was applied (<u>Fishburn et al., 2019</u>). To remove systemic signals contaminating the brain activity, short-separation regression was used: short-channel data was subtracted from the standard long-channel data (<u>Zhou et al., 2020</u>). Beer-Lambert Law was applied to transform optical density into oxygenated (HbO) and deoxygenated (HbR) hemoglobin concentration changes with a partial pathlength factor of six. Data were filtered using a third-order zero-phase Butterworth bandpass filter with cutoff frequencies of 0.01 to 0.5 Hz to remove instrumental and physiological noise.

### 2.7. Data analysis

The data of 9 participants were set aside either because they were incomplete (i.e., participants had not entirely filled the questionnaires or the *f*NIRS signal was not recorded due to a technical problem) or of insufficient quality (i.e., the *f*NIRS was too noisy to be analyzed). The data were analyzed as a function of the type of dilemma and the type of argument. The data were also analyzed based on the valence of the arguments (i.e., whether the arguments provided by the AI-advisors contradicted or validated the choice made by participants in the baseline run). In this second type of analysis, the type of dilemma factor was not included, due to the variable proportion of validating and contradicting arguments among each class of dilemmas.

#### 2.7.1. Behavioral data

Due to the binary nature of decisions, the non-normality of the data and the repeated-measures design of the study, we chose to use Generalized Estimating Equation (GEE) models to analyze participants’ decisions, change of minds and delegations. Response times were log-transformed due to their non-normal distribution and analyzed using ANOVA or dependent t-tests.

##### Decisions

A 2 x 3 [Dilemma Type (utilitarian, deontological) x Argument Type (no argument, utilitarian argument, deontological argument)] binary logistic regression was conducted on participants’ decisions (with 0 = deontological choice and 1 = utilitarian choice). A manual stepwise analysis was performed to remove non-significant interactions from the model and pairwise comparisons were carried out to further examine significant effects (α < .05).

##### Changes of mind

Four analyses were conducted to investigate participants’ changes of minds between the baseline run and the AI-advised run. A 2 x 2 [Dilemma Type (utilitarian, deontological) x Argument Type (utilitarian argument, deontological argument)] binary logistic regression was conducted on participants’ decisions to change their mind (with 0 = no change of mind and 1 = change of mind), without discriminating the nature of the change of mind (i.e., from a deontological choice to a utilitarian one and the opposite). Two supplementary 2 x 2 [Dilemma Type (utilitarian, deontological) x Argument Type (utilitarian argument, deontological argument)] binary logistic regressions were conducted on participants’ decisions to change their mind 1) from the utilitarian option to the deontological option and 2) from the deontological option to utilitarian option in response to the AI-advisors’ arguments (with 0 = no change of mind and 1 = change of mind). Finally, a 2 [Argument Valence (validating, contradicting)] binary logistic regression was conducted on participants’ decisions to change their mind (with 0 = no change of mind and 1 = change of mind).

##### Delegations versus active decisions

A 2 x 2 [Dilemma Type (utilitarian, deontological) x Argument Type (utilitarian, deontological)] and a 2 [Argument Valence (validating, contradicting)] binary logistic regressions were performed on active decisions (with 0 = delegation and 1 = active decision). Manual stepwise analyses were performed to remove non-significant interactions from all the models and pairwise comparisons were carried out to further examine significant effects (α < .05).

##### Response times

Because participants made their decision after reading the dilemmas in the baseline run versus after reading the advice of AI-advisors in the AI-advised run, the response times of each run were analyzed separately. A 2 [Dilemma Type (utilitarian, deontological)] dependent t-test was performed on the log-transformed response times when no arguments were provided to the participants (i.e., baseline run). Two analyses were performed on the response times measured in the AI-advised run. A 2 x 2 [Dilemma Type (utilitarian, deontological) x Argument Type (utilitarian argument, deontological argument)] ANOVA and a 2 [Argument Valence (contradicting, validating)] paired t-test were performed on the log-transformed response times measured in the AI-advised run. Tukey post-hoc tests were carried out to further examine significant effects (α < .05) of the ANOVA analysis.

#### 2.7.2. Subjective data

To investigate whether participants’ empathy level predicted participants’ utilitarianism rate, we conducted a multiple regression analysis with participants’ utilitarianism rate as dependent variable and the IRI subscales ratings (i.e., perspective taking, empathic concern and personal distress subscales) as predictors. A second and a third multiple regression analysis was conducted to investigate whether participants’ attitudes toward AI, empathy level and tendency to comply predicted participants’ propensity to 1) change their mind and to 2) delegate the execution of the decision, when the arguments of AI-advisors contradicted versus validated participants decisions in the baseline run, with participants’ change of mind rate as dependent variable and the ratings of IRI subscales, the NARS subscales (i.e., interaction, social and emotion) and the Gudjonsson’s compliance scale as predictors. Manual stepwise analyses were performed to remove the effects with a *p* > .15 from the models (<u>Field, 2013</u>).

#### 2.7.3. *f*NIRS data

The generalized linear model (GLM) approach was used to quantify the amplitude of evoked hemodynamic responses per ROI and Condition. The GLM was fit to the long-channel data and included all principal components of short-detector channels to account for extracerebral and physiological signal components (<u>Gagnon et al., 2014</u>). The design matrix for the GLM was generated by convolving a boxcar function at each event-onset time with the SPM canonical hemodynamic response function (<u>Abraham et al., 2014</u>). The GLM was performed with a lag-1 autoregressive noise model, to account for the correlated nature of the *f*NIRS signal components. Individual beta estimates were then averaged for each ROI, weighted by the standard error. As the oxyHb response is considered as a more sensitive indicator of changes in regional blood flow (<u>Yücel, 2021</u>), we chose to focus on the variations in oxyHb level. The use of ROIs as a factor may lead to unintended statistical bias due to the optical properties that may differ systematically between ROIs (<u>Herold et al., 2018</u>).

##### Dilemma

For the following analyses, the duration of boxcar function was 15 seconds and drift orders accounting for signal components up to 0.033 Hz were included as regression factors. To compare the hemodynamic responses observed for utilitarian dilemmas and deontological dilemmas, three 2 [Argument Valence (contradicting, validating)] paired t-test were performed on β values extracted from GLM separately for each ROI. To test whether the hemodynamic responses observed for utilitarian dilemmas and deontological dilemmas varied between the second and the third presentation, three 2 x 2 [Dilemma Type (utilitarian, deontological) x Presentation Order (second time, third time)] ANOVA were performed on β values extracted from GLM separately for each ROI. Tukey post-hoc tests were carried out to further examine significant effects (α < .05) of the ANOVA analysis.

##### Dilemma & Argument

Due to the important variability of the measurements performed in the literature (e.g., <u>Balconi & Fronda, 2020</u>; <u>Dashtestani et al., 2018</u>, <u>2019</u>; <u>Greene et al., 2004</u>; <u>Lee & Yun, 2017</u>; <u>Strait & Scheutz, 2014</u>), the hemodynamic responses observed as a function of both dilemma and argument types were assessed according to three temporal periods: 1) time-locked to the presentation of the argument, 2) time-locked 15 seconds before the response and 3) in the [-8s ; 8s] interval around the response. The duration of the boxcar function depended on the temporal periods: 30 seconds for the first, 15 seconds for the second and 16 seconds. Drift orders included as regression factors were also designed based on the temporal periods, with drift orders accounting for signal components up to 0.016 Hz for the first, 0.033 Hz for the second and 0.031 Hz for the third temporal period. For each of these three measures, three 2 x 2 [Dilemma Type (utilitarian, deontological) x Argument Type (utilitarian argument, deontological argument)] ANOVAs were performed on β values extracted from GLM, on each of the three ROIs (i.e., nine analyses in total). Tukey post-hoc tests were carried out to further examine significant effects (α < .05) of the ANOVA analyses. *Argument valence.* The hemodynamic responses observed as a function of the argument valence were also assessed according to three temporal periods: 1) time-locked to the presentation of the argument, 2) time-locked 15 seconds before the response and 3) in the [-8s; 8s] interval around the response. The duration of the boxcar function depended on the temporal periods: 30 seconds for the first, 15 seconds for the second and 16 seconds. Drift orders included as regression factors were also designed based on the temporal periods: drift orders accounting for signal components up to 0.033 Hz for the first, 0.063 Hz for the second and 0.031 Hz for the third). For each of these three measures, three 2 [Argument Valence (contradicting, validating)] paired t-tests were performed on β values extracted from GLM, on each of the three ROIs (i.e., nine analyses in total).

## 3. RESULTS

### 3.1. Behavioral results

#### 3.1.1. Decisions

The 2 x 3 [Dilemma Type (utilitarian, deontological) x Argument Type (no argument, utilitarian argument, deontological argument)] binary logistic regression conducted on decisions revealed a significant main effect of dilemma type [*B* (SE) = −1.502 (.186), CI _(95%)_ = (−1.867, − 1.137), Wald χ*²* (1) = 65.203, *p* < .001; see Figure 3.A.], with utilitarian dilemmas predicting for a high rate of utilitarian decisions (*M* = 69.41 %, *SD* = 26.85) than deontological dilemmas (*M* = 33.81 %, *SD* = 24.35). The analysis also revealed a main effect of the argument type [utilitarian argument: *B* (SE) = − .094 (.109), CI _(95%)_ = (− .307, .120), Wald χ*²* (1) = .737, *p* = .391; deontologist argument*: B* (SE) = .286 (.129), CI _(95%)_ = (.034, .538), Wald χ*²* (1) = 4.930, *p* < .05; with no argument as dummy], with significantly lower utilitarian decision rate when AI-advisors provided a deontological argument (*M* = 46.59 %, *SD* = 32.80) than a utilitarian argument (*M* = 55.02 %, *SD* = 31.00, *p* < .01) and when no argument was provided (*M* = 53.23 %, *SD* = 29.51, *p* < .05; see Figure 3.B.). No difference in utilitarian decision rate was found when no argument was provided and when a utilitarian argument was provided by the AI-advisor (*p* = .389).

**Figure 3.**
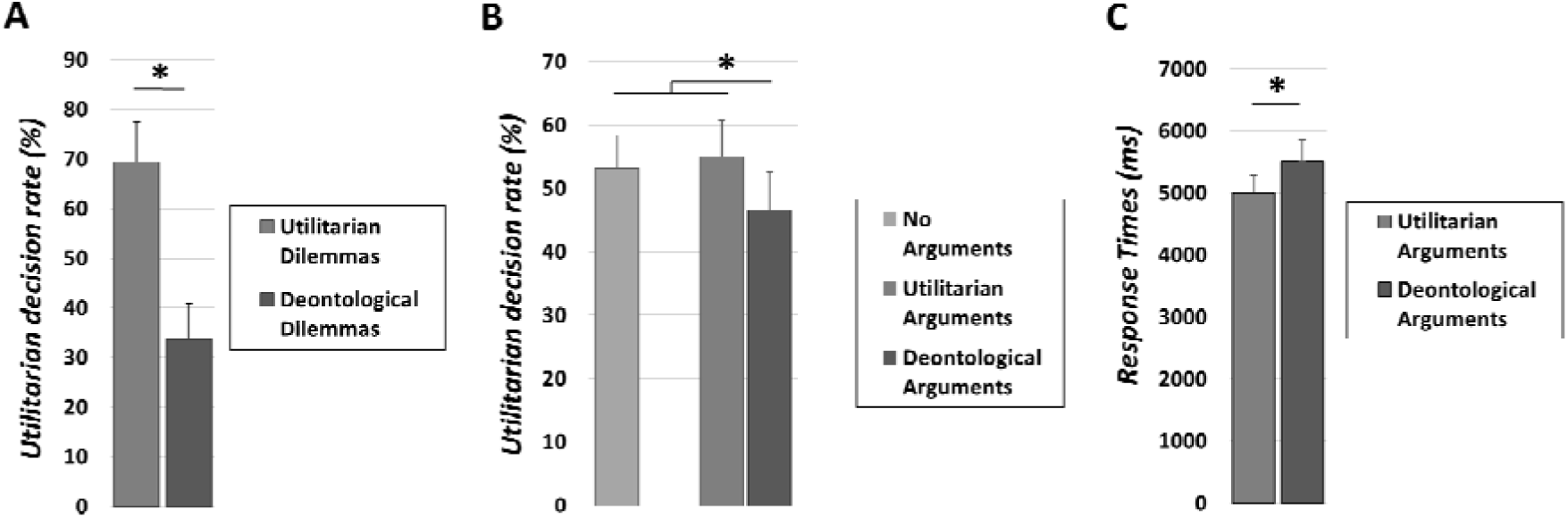
Illustration of A) the utilitarian decision rate as a function of the dilemmas type (with utilitarian dilemmas in mid-grey and deontological dilemmas in dark gray), B) the argument type (with (with no argument in light grey, utilitarian arguments in mid-grey and deontological arguments in dark grey); and C) response times as a function of the argument type in the AI-advised run.

#### 3.1.2. Changes of mind between the baseline run and the AI-advised run

##### Changes of mind as a function of the argument type

The analysis revealed that no difference in change of mind rate was found as a function of the dilemma type [*B* (SE) = .221 (.310), CI _(95%)_ = (−.386, .828), Wald χ*²* (1) = .508, *p* = .476] and the argument type [*B* (SE) = .060 (.131), CI _(95%)_ = (−.196, .316), Wald χ*²* (1) = .211, *p* = .646].

##### Changes of mind from the utilitarian to the deontological option

The analysis revealed a significant main effect of argument type [*B* (SE) = .808 (.256), CI _(95%)_ = (.307, 1.309), Wald χ*²* (1) = 9.987, *p* < .01; see Figure 4.A], with deontological arguments (*M* = 8.42 %, *SD* = 11.82) predicting for a greater amount of changes of mind from a utilitarian to a deontological decision than utilitarian arguments (*M* = 3.94 %, *SD* = 5.59). The main effect of dilemma type [*B* (SE) = .282 (.293), CI _(95%)_ = (−.292, .856), Wald χ*²* (1) = .926, *p* = .336] did not reach significance.

**Figure 4.**
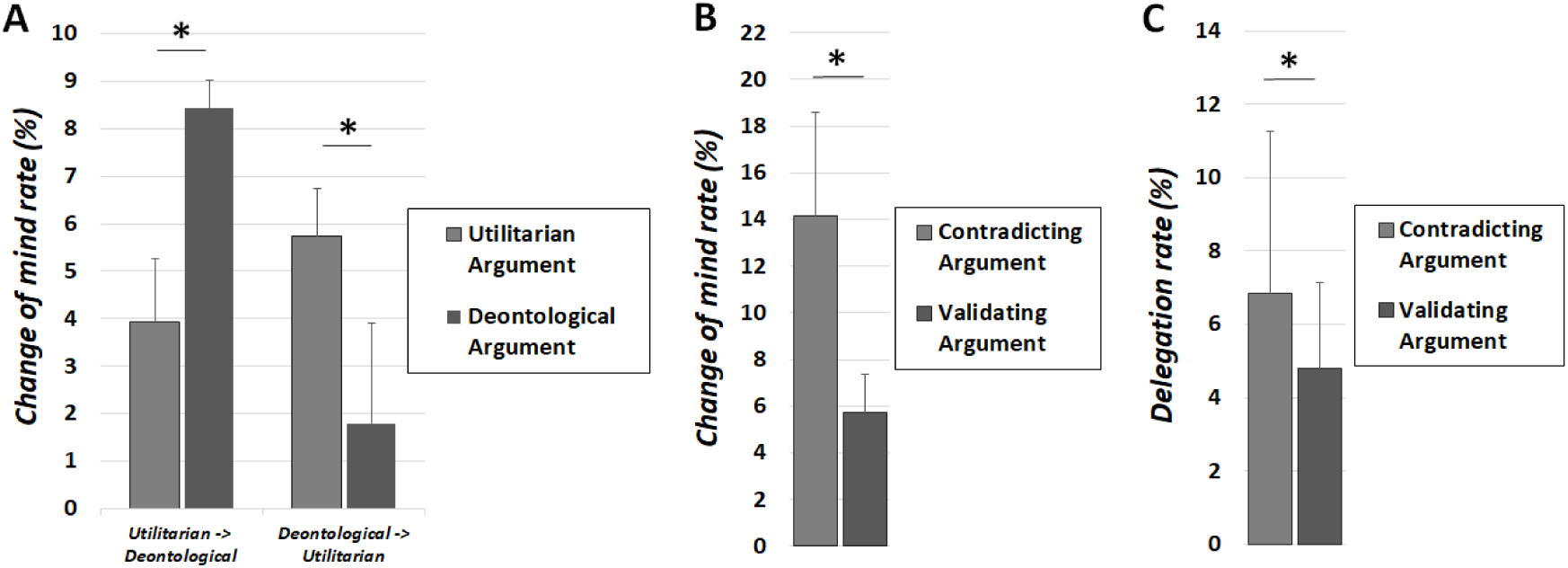
Illustration of A) the change of mind rate as a function of the argument type from deontological option to utilitarian option (left) and from utilitarian option to deontological option, B) the change of mind rates and C) the delegation rates as a function of the valence of the arguments.

##### Changes of mind from the deontological to the utilitarian option

The analysis revealed a significant main effect of argument type [*B* (SE) = −1.204 (.344), CI _(95%)_ = (−1.877, −.531), Wald χ*²* (1) = 12.291, *p* < .001; see Figure 4.A], with utilitarian arguments predicting for a greater amount of changes of mind from the deontological to the utilitarian option (*M* = 5.73 %, *SD* = 7.38) than deontological arguments (*M* = 1.79 %, *SD* = 3.33). The main effect of dilemma type [*B* (SE) = .100 (.436), CI _(95%)_ = (−.755, .955), Wald χ*²* (1) = .053, *p* = .818] did not reach significance.

##### Changes of mind as a function of the argument valence

In the AI-advised run, 17 participants showed a greater change of mind rate in response to the contradicting arguments than the validating arguments, 10 participants showed the same amount of change of minds in response to the validating and the contradicting arguments, and 5 participants never changed their minds at all. The analysis revealed a main effect of argument valence [*B* (SE) = − .967 (251), CI _(95%)_ = (− 1.469, − .483), Wald χ*²* (1) = 15.080, *p* < .001; see Figure 4.B], with contradicting arguments predicting for a greater change of mind rate (*M* = 14.16 %, *SD* = 19.93) than validating arguments (*M* = 5.73 %, *SD* = 9.16).

#### 3.1.3. Delegations versus active decisions in the AI-advised run

In the AI-advised run, 14 participants never delegated the execution of the decision to the AI-advisors, while 17 participants made at least one delegation. The results revealed a rather low mean delegation rate (*M* = 5.83 %, *SD* = 14.13).

##### Active decisions as a function of the argument type

The analysis performed on active decisions revealed no significant main effects of dilemma type effect [*B* (SE) = −.223 (.230), CI _(95%)_ = (− 675, .229), Wald χ*²* (1) = .936, *p* = .333] or argument type [*B* (SE) = .032 (.183), CI _(95%)_ = (− .327, .390), Wald χ*²* (1) = .030, *p* = .862].

##### Active decisions as a function of the valence of the argument

The analysis performed on active decisions revealed a significant main effect of argument valence [*B* (SE) = .417 (.186), CI _(95%)_ = (.053, .781), Wald χ*²* (1) = 5.029, *p* < .05], with significantly fewer active decisions when the AI-advisors provided contradicting arguments (*M* = 93.15 %, *SD* = 15.29) than validating arguments (*M* = 95.19 %, *SD* = 13.36; see Figure 4.C).

#### 3.1.4. Response times

##### Response times in the baseline run

No difference in response times was found in response to utilitarian and deontological dilemmas [*t* (31) = − 1.260, *p* = .217, CI*_95%_* = (− .135; .032)] when no arguments were provided by the AI-advisors (i.e., baseline run).

##### Response times as a function of the argument type

The analysis revealed a main effect of arguments type [*F* (1, 30) = 8.708, *p* < .01, η*p²* = .225], with longer response times when AI-advisors provided deontological arguments (*M* = 5515 ms, *SD* = 1477) than utilitarian arguments (*M* = 5008 ms, *SD* = 1291; see Figure 3.C). The main effect of dilemma type [*F* (1, 30) = .356, *p* = .555, η*p²* = .012] and Dilemma Type x Argument Type interaction [*F* (1, 30) = .635, *p* = .432, η*p²* = .021] did not reach significance.

##### Response times as a function of the argument valence

No difference in response times was found as a function of the argument valence [*t* (31) = − 0.223, *p* = .825, CI*_95%_* = (− .021; .017)].

#### 3.1.5. Subjective & behavioral data

Participants’ level of empathy was not predictive of their utilitarian decision rate in the first phase of the experiment [*F* (1, 29) = .291, *p* = .594, R^2^ = .010, R^2^_adjusted_ = − .024; see Table 1 for a summary of the model]. However, in the second phase of the experiment participants’ tendency to comply and tendency to experience positive emotions toward robots/A.I. significantly were predictive of participants’ propensity both to change their mind [*F* (3, 27) = 7.790, *p* < .001, R^2^ = .464, R^2^_adjusted_ = .404] and to delegate the execution of the action [*F* (3, 27) = 4.812, *p* < .01, R^2^ = .348, R^2^ = .276] when being contradicted (versus validating) by the AI-advisors (see Table 1 for a summary of the model).

**Table 1.** Summary of the multiple regression analyses performed on the questionnaire ratings and the decision rates.

| Multiple Regression Analysis | Unstandardized Coefficients |  | Standardized Coefficients | t | p | CI (95%) |  |
| --- | --- | --- | --- | --- | --- | --- | --- |
|  | B | SE | Beta |  |  | Inferior | Superior |
| Utilitarianism in the baseline run |  |  |  |  |  |  |  |
| Constant | .362 | .313 |  | 1.157 | .257 | -.278 | 1.002 |
| IRI_Perspective Taking subscale | .036 | .066 | .100 | .539 | .594 | -.100 | .172 |
| Propensity to change mind |  |  |  |  |  |  |  |
| Constant | - 72.620 | 23.639 |  | - 3.072 | .005 | -121.122 | - 24.117 |
| Gudjonsson's compliance scale | 11.892 | 2.830 | .597 | 4.202 | .000* | 6.036 | 17.685 |
| NARS_Emotion subscale | 3.545 | 1.429 | .355 | 2.492 | .020* | .613 | 6.477 |
| IRI_Perspective Taking subscale | 8.630 | 4.306 | .284 | 2.057 | .055 | -.205 | 17.464 |
| Propensity to delegate |  |  |  |  |  |  |  |
| Constant | -60.502 | 23.84 |  | -2.538 | .017 | -109.419 | -11.586 |
| Gudjonsson's compliance scale | 8.318 | 2.863 | .457 | 2.905 | .007* | 2.444 | 14.193 |
| NARS_Emotion subscale | 3.523 | 1.441 | .385 | 2.445 | .021* | .566 | 6.479 |
| IRI_Perspective Taking subscale | 7.56 | 4.342 | .272 | 1.741 | .093 | -1.35 | 16.47 |

### 3.2. *f*NIRS results

The 2 x 2 [Dilemma Type (utilitarian, deontological) x Argument Type (utilitarian argument, deontological argument)] ANOVA performed on the hemodynamic responses time-locked to the presentation of the arguments and measured at the right ROI was the only analysis that reached significance. For the sake of clarity, non-significant analyses are reported in Table 2.

**Table 2.** Summary of the statistical analyses performed on the *f*NIRS measurements.

| **ANOVA** | **Left ROI** |  |  | **Rostral ROI** |  |  | **Right ROI** |  |  |
| --- | --- | --- | --- | --- | --- | --- | --- | --- | --- |
|  | *F(1,30)* | *p* | *ηp<sup>2</sup>* | *F(1,30)* | *p* | *ηp<sup>2</sup>* | *F(1,30)* | *p* | *ηp<sup>2</sup>* |
| **Dilemma x Presentation Order (2nd vs 3rd)** |  |  |  |  |  |  |  |  |  |
| Dilemma | .120 | .732 | .004 | 1.194 | .283 | .038 | .481 | .493 | .016 |
| Order | .231 | .634 | .008 | .013 | .909 | .000 | .027 | .869 | .001 |
| Dilemma x Order | .256 | .616 | .008 | **4.668** | **.039\*** | **.135** | .013 | .910 | .000 |
| **Dilemma x Argument (Time Locked to Argument)** |  |  |  |  |  |  |  |  |  |
| Dilemma | 2.387 | .133 | .007 | 1.562 | .221 | .049 | .006 | .941 | .000 |
| Argument | .348 | .560 | .011 | 1.078 | .308 | .035 | **8.109** | **.007\*** | **.213** |
| Dilemma x Argument | .037 | .849 | .001 | .061 | .806 | .002 | 1.749 | .196 | .055 |
| **Dilemma x Argument (Before Answer)** |  |  |  |  |  |  |  |  |  |
| Dilemma | .025 | .876 | .001 | .180 | .674 | .006 | .071 | .792 | .002 |
| Argument | .270 | .607 | .009 | .179 | .286 | .038 | .020 | .888 | .001 |
| Dilemma x Argument | 1.955 | .172 | .061 | .208 | .651 | .007 | .383 | .546 | .013 |
| **Dilemma x Argument (Around answer)** |  |  |  |  |  |  |  |  |  |
| Dilemma | 1.441 | .239 | .046 | 2.605 | .117 | .080 | .420 | .522 | .014 |
| Argument | .001 | .969 | .000 | 2.977 | .095 | .090 | 1.017 | .321 | .033 |
| Dilemma x Argument | 2.371 | .134 | .073 | 1.904 | .178 | .060 | .000 | .982 | .000 |

| **Paired t-tests** | **Left ROI** |  |  | **Rostral ROI** |  |  | **Right ROI** |  |  |
| --- | --- | --- | --- | --- | --- | --- | --- | --- | --- |
|  | *t* | *df* | *p* | *t* | *df* | *p* | *t* | *df* | *p* |
| **Dilemma** |  |  |  |  |  |  |  |  |  |
| Time-Locked to Dilemma | .336 | 30 | .739 | .012 | 30 | .192 | -.186 | 30 | .854 |
| **Argument Nature** |  |  |  |  |  |  |  |  |  |
| Time-Locked to Argument | 1.232 | 30 | .228 | 1.064 | 30 | .296 | -.045 | 30 | .964 |
| Before Response | -1.330 | 30 | .193 | .967 | 30 | .341 | -.277 | 30 | .783 |
| Around Response | -.043 | 30 | .966 | -.584 | 30 | .563 | -.045 | 30 | .964 |

#### Hemodynamic response time-locked to the argument

The ANOVA analysis conducted on the right ROI revealed a significant main effect of argument type [*F*(1, 30) = 8.109, *p* < .01, *np² =* .213, see Figure 5], with a higher consumption of oxygenated hemoglobin observed in response to utilitarian arguments (*M* = .019, *SD* = .010) than to deontological arguments (*M* = − .013, *SD* = .013). The main effect of dilemma type [*F*(1, 30) = .006, *p* = .941, *np² =* .000] and the Dilemma Type x Argument Type interaction [*F*(1, 30) = 1.749, *p* = .196, *np² =* .055] did not reach significance.

**Figure 5.**
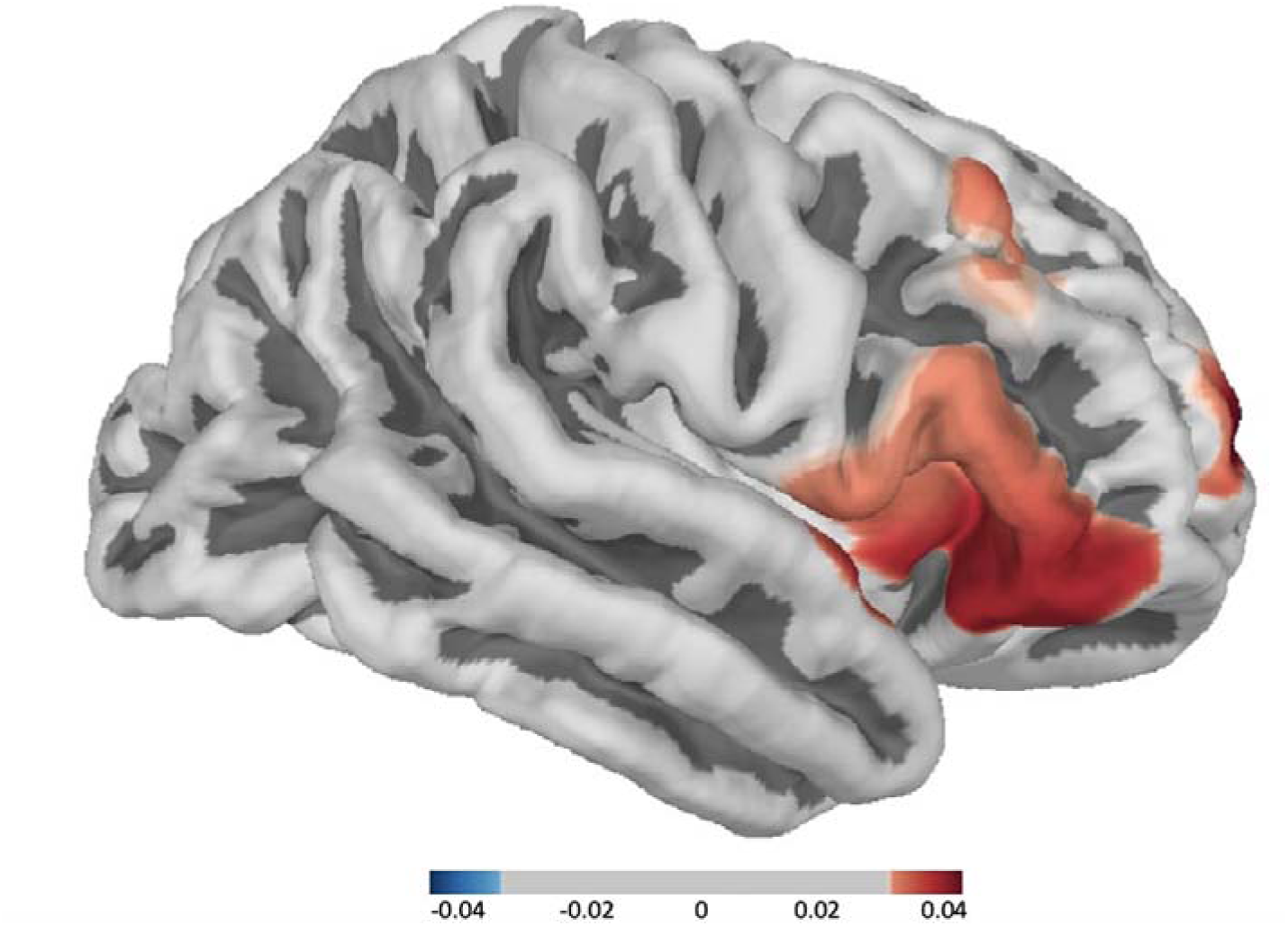
Topographic plot of the contrast between the responses to utilitarian arguments > deontological arguments.

## 4. DISCUSSION

The present experiment aimed at investigating the influence of AI-advisors on human moral decision-making and the associated brain correlates. Participants were presented with sacrificial dilemmas and were tasked with making moral decisions either independently themselves (i.e., baseline run) or with AI-advisors (i.e., AI-advised run). Their brain activity was assessed during the AI-advised run using an *f*NIRS system.

### 4.1. Deciding on utilitarian and deontological dilemmas

In both the baseline and the AI-advised runs, participants were significantly more likely to choose the utilitarian option in response to utilitarian dilemmas (i.e., ∼ 70%) and the deontological option in response to deontological dilemmas (i.e., ∼ 67%). This result serves as a manipulation check confirming that the two types of dilemmas were properly defined and balanced based on the first rating study. In the AI-advised run, no modulation of DLPFC activity was observed as a function of the type of dilemma (not in line with <u>Greene et al., 2001</u>, <u>2004</u>). It is possible that this lack of difference in brain activity stems from the fact that all dilemmas had already been presented during the baseline run. Participants may have engaged in less thorough analysis of the dilemmas when they were presented for the second and third time during the AI-advised run, resulting in an absence of brain activity difference. In contradiction with some previous studies (<u>Gleichgerrcht & Young, 2013</u>; <u>Nasello et al., 2021</u>; <u>Takamatsu, 2018</u>), participants’ self-assessed level of empathy was not predictive of their level of utilitarianism. This lack of association between utilitarianism and empathy in the current study is not unexpected. A recent meta-analysis demonstrated that the role of empathy in moral judgments can strongly vary, oscillating between a small/moderate to limited/insignificant link depending on the moral decisions (<u>Nasello & Triffaux, 2023</u>).

### 4.2. Change of mind in response to AI-advisors’ arguments

Overall, participants were significantly more likely to change their mind 1) from the utilitarian to the deontological option in response to deontological arguments than to utilitarian ones and 2) from the deontological to the utilitarian option in response to utilitarian arguments than to deontological ones. Moreover, participants changed significantly more their mind when the arguments of AI-advisors contradicted than validated the decisions they made in the baseline run. Taken together, these results demonstrate that AI-advisors can have a significant influence on moral decision-making (in line with <u>Bai et al, 2023</u>; <u>Hertz & Weise; 2018</u>; <u>Köbis et al., 2021</u>; <u>Leib et al., 2024</u>). While 17 participants were more likely to change their minds in response to contradicting arguments than to validating ones, 10 participants showed an equal change of mind rate in response to both types of arguments, and 5 participants never changed their minds at all. Therefore, despite the statistical significance of these findings, it is important to note that the AI advisors’ arguments did not affect all participants uniformly.

Interestingly, participants’ propensity to change their mind when contradicted by AI- advisors was predicted both by their self-estimated tendency to comply (i.e., Gudjonsson’s compliance scale) and their disposition to experience negative emotions toward non-human agents (i.e., inverted emotion NARS subscale). In other words, the influence of AI-advisors appears to depend on human advisees’ susceptibility to obey and to fear non-human agents, as previously found with human advisors (e.g., <u>Fabre et al., 2022</u>).

### 4.3. Impact of the utilitarian versus deontological arguments of AI-advisors

Being advised to make deontological choices by the AI-advisors seems to have prompted participants to thoroughly reassess their decisions, as indicated by the longer response times observed in response to deontological arguments than to consequentialist ones. Interestingly, greater activity of right-DLPFC was also measured in response to utilitarian arguments than to deontological ones. In various previous studies, increased activity in the right-DLPFC was found to predict for (more) utilitarian decisions (e.g., <u>Greene et al., 2004</u>; <u>Jeurissen et al., 2014</u>; <u>Lee & Yun, 2017</u>; <u>Patil et al., 2021; Zheng et al., 2018</u>). The right-DLPFC is thought to underpin the cognitive control processes necessary to appraise the emotional response to moral dilemmas (<u>Greene et al., 2004</u>; <u>Tassy et al., 2012</u>). Being provided with deontological arguments by the AI-advisors appears to have decreased the appraisal of the affective response to the sacrificial moral dilemmas, ultimately resulting in a significant decrease in participants’ utilitarianism.

Human beings are acutely aware that the moral decisions they make shape both their self-image and the way they are perceived by others (<u>Bauman & Helzer, 2023</u>; <u>Brambilla et al., 2021</u>; <u>Carlson & Furr, 2009</u>; <u>Carlson et al., 2011; Macko, 2020</u>; <u>Plaks et al., 2021</u>; <u>Reynolds et al., 2019</u>; <u>Rom & Conway, 2018; Uhlmann et al., 2015</u>). As a result, they tend to adjust both their moral choices and their advice to align with the way they wish to present themselves (<u>Jin & Peng, 2021</u>; <u>Macko, 2020</u>; <u>Polman & Ruttan, 2022; Rom & Conway, 2018; Trémolière & Rateau, 2023</u>). Compared to those who behave in a deontological way, individuals who make utilitarian decisions are generally viewed as more competent, but less warm, trustworthy, empathetic, moral, prosocial, and attractive (<u>Brown & Sacco, 2019</u>; <u>Capraro et al., 2018</u>; <u>Everett et al., 2018</u>; <u>Rom et al., 2017</u>; <u>Rom & Conway, 2018</u>; <u>Sacco et al., 2017</u>; <u>Uhlmann et al., 2013</u>). Moral character being far more influential than competence when forming impressions (<u>Brambilla et al., 2011</u>, <u>2012</u>; <u>Goodwin et al., 2015</u>; <u>Leach et al., 2007</u>), human beings tend to adapt their moral preferences to appear more deontological than they actually are (e.g., <u>Bostyn & Roets, 2017; Lee et al., 2018; Reynolds et al., 2019; Sacco et al., 2017)</u>, as a strategical move to avoid the personal and social costs associated with utilitarianism (<u>Brown & Sacco, 2019</u>; <u>Everett et al., 2018</u>; <u>Goldstein-Greenwood et al., 2020</u>; <u>Rom et al., 2017</u>; <u>Szekely & Miu, 2014</u>; <u>Uhlmann et al., 2013</u>). Compared to non-human agents, human beings are less strongly expected by their peers to make utilitarian choices and are typically judged more negatively when they do, particularly in deontological dilemma situations (e.g., <u>Chu & Liu, 2023</u>; <u>Malle et al., 2015; Zhang et al., 2022</u>). Therefore, making a utilitarian decision after receiving a deontological argument from an AI-advisor exposed participants to both the personal and/or social costs of acting in a utilitarian manner (e.g., <u>Reynolds et al., 2019; Sacco et al., 2017)</u> and those of seeing themselves and/or being perceived by others as more cold-blooded than machines (<u>Chu & Liu, 2023</u>). We hypothesize that the reevaluation process that occurred when the AI-advisors provided deontological arguments reflects participants’ motivation to preserve their (self-)image (<u>Bostyn & Roets, 2017</u>; <u>Macko, 2020</u>; <u>Reynolds et al., 2019</u>; <u>Rom & Conway, 2018</u>).

The present results contradict Greene’s *moral dual-process theory*, which posits that deontological decisions mainly result from a fast, automatic and emotional decision process underpinned by System 1 (<u>Greene, 2015</u>). However, they align with Moll and colleagues’ view, asserting that cognition and emotion are continuously integrated during moral decision-making and that motivational power c here supposedly preserving one’s (self-)image c can lead to a voluntarily down- or up-regulation of affective reactions (<u>Moll et al., 2005, 2008; Ochsner et al., 2004</u>).

### 4.4. Delegating the execution of the action to the AI-advisors

In the AI-advised run, participants were given the possibility to delegate the execution of the decision to the AI-advisors by not replying for 15 seconds. On average, participants chose to delegate about 6% of the time. The results revealed significant variability among participants in their willingness to delegate the decision execution to the AI-advisors, with 14 participants never opting to delegate, and 17 participants doing it at least once. During the post-experimental debriefing, all 14 participants who never chose to delegate reported feeling uncomfortable with the notion of allowing a machine to execute such actions, expressing a preference for maintaining control (in line with <u>Bigman & Gray, 2018</u>; <u>Castelo et al., 2019</u>; <u>Dietvorst et al., 2015, 2018</u>; <u>Gogoll & Uhl, 2018</u>). Conversely, the 17 participants who delegated the execution of actions at least once reported that certain decisions were so challenging to execute that they felt relieved to be able to delegate them to the AI-advisors (in line with <u>Drugov et al., 2014</u>; <u>De Melo et al., 2016</u>; <u>Sloane & Moss, 2019</u>). A significant increase in delegation (i.e., decrease in active decisions) was observed when AI-advisors provided arguments contradicting the decisions made by participants in the baseline run. Moreover, the propensity of participants to delegate the execution to AI-advisors after being contradicted versus validated was predicted both by their self-estimated propensity to comply and tendency to experience negative emotions toward non-human agents. Based both on literature and participants’ feedback, this result suggests that participants’ decisions to delegate the execution of the actions to the AI-advisors might reflect a willingness to surrender decision-making authority to the machine to avoid the negative emotional response associated with the execution of the action (<u>Drugov et al., 2014</u>; <u>De Melo et al., 2016</u>; <u>Sloane & Moss, 2019</u>).

### 4.5. Limitations and further work

In the present study, participants were tasked with deciding on fictitious sacrificial dilemmas that may not fully represent the moral decisions individuals typically face in everyday life. Consequently, further research is warranted to assess the influence of AI moral advisors on the moral decisions humans regularly encounter in their personal and professional lives. This study is one of the first to investigate the brain activity associated with moral decision-making advised by AI-agents (<u>Goodyear et al., 2016</u>; <u>2017</u>). As previous *f*NIRS studies investigating moral decision-making (<u>Balconi & Fronda, 2020</u>; <u>Dashtestani et al., 2018</u>, <u>2019</u>; <u>Lee & Yun, 2017</u>; <u>Strait & Scheutz, 2014</u>), we focused on the activity of the PFC, a key structure involved in moral decision-making. However, the activity of various other brain structures underpinning advice taking (e.g., <u>Goodyear et al., 2016</u>, <u>2017</u>; <u>Meshi et al., 2012</u>) and/or moral judgment/decision-making (e.g., <u>Casebeer, 2003</u>; <u>Greene et al, 2001</u>, <u>2004</u>; <u>Moll et al., 2005</u>; <u>Moll & de Oliveira-Souza, 2007</u>), such as for instance the anterior cingulate cortex, the ventromedial PFC or the temporal-parietal junction, was not investigated due to the inherent constraints associated with the use of *f*NIRS (i.e., limited number of channels and measurements restricted to the outer cortex; <u>Quaresima & Ferrari, 2019</u>). Replicating the present study using *f*MRI would further improve our understanding of the involvement of both subcortical and (outer and inner) cortical structures during AI-advised moral decision-making.

Finally, further work is necessary to confirm that participants lowered their level of utilitarianism in response to the deontological arguments provided by the AI-advisors to preserve their (self-)image and to determine the extent to which this behavior is driven by self-oriented versus other-oriented motives (e.g., <u>Reynolds et al., 2019; Sacco et al., 2017)</u>.

## Conclusion

The results of the present study demonstrate that AI-agents can influence moral decision-making in human beings. They confirm the necessity to define clear rules to regulate the way artificial moral advisors are trained and used in everyday life (e.g., <u>Constantinescu et al., 2022</u>; <u>IEEE, 2017</u>; <u>Kobis et al., 2019</u>; <u>Russell et al., 2015</u>). Some authors have proposed to adjust the “behavior” of artificial moral advisors to the moral preferences of the user (e.g., <u>Giubilini & Savulescu, 2018</u>), which appears as a good solution at least for a personal use of this technology. Given the influence of artificial moral advisors, particular attention should be given to their use in a professional context c especially in the military (<u>Svenmarck et al., 2018</u>) c in which the moral preferences of the users might not align with those of the organization they work for (<u>IEEE, 2017</u>). Further research is therefore necessary to define the rules for proper and ethical use of artificial moral advisors (<u>Irving & Askell, 2019</u>).

