## Supplementary for "Making Moral Decisions With Artificial Agents As Advisors. An *f*NIRS Study"

**Rating Study 1: Selection of utilitarian and deontological dilemmas**

A first rating study was conducted to select the moral dilemmas of the main experiment. We aimed at selecting an equal number of utilitarian dilemmas and deontological dilemmas, the two groups of dilemmas being balanced in terms of choice ratio (e.g., a ~ 80% utilitarian choice dilemma had to be balanced with an ~ 80% deontological choice dilemma). We analyzed the literature and pre-selected, translated and/or adapted 24 moral dilemmas that had been used in previous studies (e.g., Foot, 1978; Grassian, 1981; Greene et al., 2001; Manfrinati et al., 2013).

A website hosted by the University of Toulouse was created to conduct the on-line rating study. The link to the study was shared on various social media (i.e., LinkedIn, Facebook). When clicking on the link, participants were directed to the welcome page of the website. They were informed that the experiment would not last more than ten minutes and that they should click on the “participate” icon to start the experiment. After they clicked on the icon, they were redirected to the second page. They were explained that their IP address would not be collected to guarantee the respect of their anonymity and kindly requested to participate in the experiment only once. They were informed that they were free to leave the experiment at any time (i.e., incomplete responses would not be accounted for) and that once they had clicked on the “finalize” icon at the end of the experiment, they would no longer have the possibility to modify or delete their answers. By clicking a second time on the icon “participate” they acknowledged that they had read and understood the instructions and accepted the terms of the experiment. Participants were then presented with eight (randomly selected) of the 24 pre-selected dilemmas. Their task was to decide between a *utilitarian option* consisting in maximizing the number of survivors and a *deontological option* consisting in avoiding causing harm to individuals unrelated to the situation and not intervene. After deciding on the eight dilemmas, they were asked to indicate their age and gender information and to click on the icon “Finalize” to validate their participation in the study. Finally, they were thanked and encouraged to share the link of the study on social media.

226 participants (157 males, 64 females and 4 non-informed) took part in this on-line rating study. Each dilemma was evaluated by 74 to 77 participants. Based on the results of this rating study, nine deontological and nine utilitarian dilemmas, with respectively comparable deontological/utilitarian and utilitarian/deontological choice ratios, were selected among the 24 pre-selected moral dilemmas that were evaluated in the study. An independent t-test analysis [*t* (16) = .191, *p* = .851, CI*_95%_* = (-8.234; 9.867)] revealed that the mean utilitarian choice rate observed for the nine utilitarian dilemmas (*M* = 69.92 %, *SD* = 9.77) was not significantly different from the mean deontological choice rate observed for the nine deontological dilemmas (*M* = 70.74%, *SD* = 8.28), confirming that the two groups of dilemmas were properly balanced. Both the French and the English version of the 18 dilemmas that were used in the present study are presented in the Supplementary materials (see Table 1S).

**Rating Study 2: Selection of the utilitarian and deontological arguments**

A second rating study was conducted (concomitantly to the rating study presented in the previous subsection) to select one utilitarian argument and one deontological argument per dilemma (i.e., respectively arguing that *“it is necessary to sacrifice the few to save the many”* versus that *“unrelated people should not be sacrificed to save others”*) ‒ the two classes of arguments being balanced in terms of persuasiveness. The procedure of this rating study was identical to the one previously described. Participants were presented with eight (randomly selected) of the 24 pre-selected dilemmas. Half of the dilemmas were accompanied by three to four utilitarian arguments and the other half were accompanied by three to four deontological arguments. Participants were asked to evaluate the level of persuasiveness of each argument on a 7-point Likert scale (from 1 = *not convincing at all* to 7 = *extremely convincing*).

356 participants (229 males, 106 females and 21 non-informed) took part in this second rating study. Each of the 162 arguments was evaluated by 51 to 63 participants (*M* = 56.83, *SD* = 2.84). Based on the results, we selected one utilitarian argument and one deontological argument for each dilemma, characterized by a similar level of persuasiveness. A 2 x 2 (Dilemma Type [utilitarian, deontological] x Argument Type [utilitarian, deontological]) ANOVA was conducted on the persuasiveness mean ratings of the arguments. The analysis revealed no significant main effects of dilemma type [*F* (1, 32) = .000, *p* = .988, *ηp²* = .000] or argument type [*F* (1, 32) = .027, *p* = .870, *ηp²* = .027], and no significant Dilemma Type x Argument Type interaction [*F* (1, 32) = 1.185, *p* = .284, *ηp²* = .036], confirming that the arguments were properly balanced in terms of persuasiveness. The two arguments (i.e., one utilitarian and one deontological) associated with each of 18 dilemmas that were used in the present study are presented both in French and in English in the Supplementary materials (see Table 1S).
